# Naturally occurring mutations in replication proteins of a small RNA virus that alter the number, sizes, and relative abundances of subgenomic RNAs

**DOI:** 10.64898/2025.12.22.695885

**Authors:** Camila Perdoncini Carvalho, Deya Wang, Junping Han, Khwannarin Khemsom, Hanqiao Chen, Yizhi Jane Tao, Feng Qu

## Abstract

Many positive-strand (+) RNA viruses produce subgenomic RNAs (sgRNAs) in infected cells. sgRNAs are synthesized by virus-encoded replication proteins (RPs), but whether RPs regulate the number and sizes of sgRNAs remains largely unknown. We report multiple naturally occurring mutations within the RPs of turnip crinkle virus (TCV) that alter the number, sizes, and relative abundances of TCV sgRNAs. TCV is a (+) RNA virus that normally produces two sgRNAs: the 1,724-nucleotide (nt) sgRNA1 expressing movement proteins, and the 1,449-nt sgRNA2 expressing capsid protein. A single amino acid change, A113V, within a region shared by TCV RPs p28 and p88, diminished sgRNA1 levels and delayed viral systemic spread. Interestingly, three second-site RP mutations emerged in infected plants that, alone or in combination with A113V, resulted in over-production of sgRNA1 or accumulation of two alternative sgRNAs of 1,876 and 1,601 nt, and rescued A113V defects. The alternative sgRNAs originated from nearly identical recombination events, their size difference reflecting varying 5’ extensions. Structural modeling of the TCV replication complex showed a conical ring architecture containing p28, p88, and a partially double-stranded RNA. While A113 may interact with TCV genomic RNA or host factors to promote (-) sgRNA1 synthesis, the second-site mutations likely influenced binding and entry of RNA template into the p88 active site. They may stall RNA synthesis at specific hotspots and stimulate template switching, thereby generating alternative (–) sgRNAs. Our findings reveal previously unrecognized constraints on viral RPs that ensure production of sgRNAs with precise sizes and abundances.

**Author summary:** Many (+)-strand viruses, such as SARS-CoV-1 and -2, Chikungunya virus, and tomato mosaic virus, synthesize subgenomic RNAs (sgRNAs) during cellular infections. Targeting sgRNA production could prove to be an effective antiviral strategy, as sgRNAs are needed to express diverse proteins critical for viral survival and transmission. Although sgRNA production requires virus-encoded replication proteins (RPs), whether RPs also dictate the number, sizes, and relative abundances of viral sgRNAs remain to be thoroughly investigated. We identified and characterized four naturally occurring mutations in RPs of the plant-infecting turnip crinkle virus that specifically perturbed one of the sgRNAs. Our findings uncover novel constraints on viral RPs that safeguard sgRNA integrity, and avail them as potential targets for controlling pathogenic viruses.

## Introduction

Viruses with single-stranded, positive sense (+) RNA genomes pose major threats to human and animal health as well as food security. An intriguing feature of many (+) RNA viruses, including the human-infecting SARS-CoV-2, Chikungunya virus, and the plant-infecting tomato mosaic virus (ToMV), is their ability to produce subgenomic RNAs (sgRNAs) during infection (1, 2). sgRNAs are typically not packaged into virions. Instead, they are synthesized inside infected cells, and serve as mRNAs for various late-expressing viral proteins (3). sgRNAs generally share identical 3’ termini with the genomic RNAs (gRNAs) of cognate viruses but possess distinct 5’ termini that map to specific nucleotide (nt) positions within gRNAs. The exact coordinates of sgRNA 5’ termini, hence the sizes of sgRNAs, are determined by secondary structures formed through intra-strand base pairing of viral gRNAs, along with conserved sequence motifs (4–7). Virus-encoded replication proteins (RPs), including auxiliary replication proteins (ARP) and RNA-dependent RNA polymerases (RdRp), are essential for sgRNA transcription (8, 9). However, whether these RPs also play a direct role in determining sgRNA number and sizes has not been thoroughly examined.

Here we report several naturally occurring mutations within the RPs of a small (+) RNA virus that alter the number, sizes, and relative abundances of sgRNAs. These mutations were identified in plant cells infected by turnip crinkle virus (TCV) (10). TCV is a plant-infecting (+) RNA virus with a 4,054-nt genome encoding five proteins (Fig. 1A). The two 5’-proximal proteins, p28 and p88, are directly translated from TCV gRNA and are essential for both gRNA replication and sgRNA transcription. The p28 ARP is produced at much higher levels than the p88 RdRp, the latter being product of infrequent translational readthrough of the p28 stop codon (Fig. 1A) (11). The three 3’-proximal proteins, p8, p9, and p38, are not translated directly from gRNA. Rather, they are translated from two sgRNAs synthesized inside infected cells. p8 and p9, two small proteins required for viral cell-to-cell movement, are translated from the 1,724-nt sgRNA1, whereas p38, the viral capsid protein and suppressor of RNA silencing-mediated host defense, is translated from the 1,449-nt sgRNA2 (Fig. 1A). Although these sgRNA-derived proteins are not required for viral genome replication in single cells, they are needed for other processes such as virus particle assembly, cell-to-cell spread, and evasion of host defenses.

**Figure 1.**
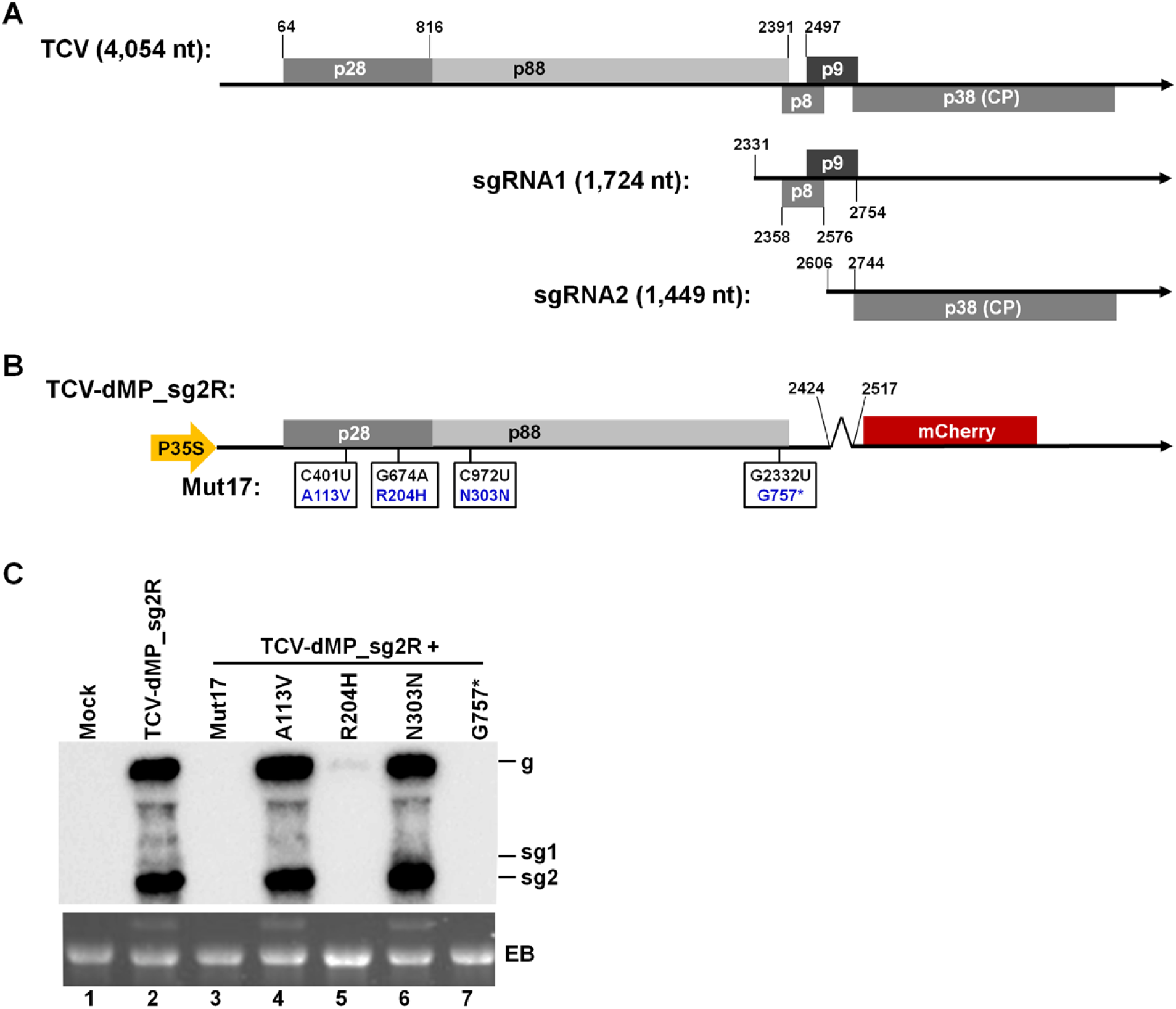
**A**. Schematic representation of the wildtype (wt) TCV genome. TCV-encoded proteins are depicted as gray boxes. The nucleotide (nt) coordinates for p28 and p88 ORFs, as well as the start of p9 ORF, are highlighted on the genomic (g) RNA diagram. Coordinates for sgRNA start positions, as well as proteins they template, are shown on the two subgenomic (sg) RNA diagrams. **B.** Schematic representation of TCV-dMP_sg2R and the four single-aa mutations that were part of the Mut17 mutant identified previously (12). **C.** Northern blot hybridization (NB) result showing the replication levels of Mut17, as well as four single-aa mutants harboring the individual Mut17 mutations. EB refers to the ethidium bromide (EB)-stained gel.

We previously identified numerous naturally occurring TCV mutations through single-cell infections (12). We hence speculated that some might compromise TCV sgRNA genesis. To test this idea, we re-examined a mutant containing four different point mutations, three of which resulted in amino acid (aa) changes in p28/p88 (Fig. 1B). Intriguingly, when introduced into TCV replicons, one mutation, C401U, causing the aa substitution A113V, drastically reduced sgRNA1 levels. An extensive analysis of plants infected by the A113V mutant identified additional TCV mutants harboring compensatory second-site mutations in p88. Some of the second-site mutations, either alone or in combination with A113V, further perturbed sgRNA1 levels and led to the production of alternative sgRNAs of distinct sizes. These findings uncover novel mechanisms by which TCV – and potentially other (+) RNA viruses – relies on intricate interactions between the gRNA and specific RP aa residues to safeguard sgRNA integrity.

## Results

### The A113V mutation within the p28 portion of p28/p88 diminishes sgRNA1 levels and delays TCV systemic spread

We previously reported the isolation of numerous TCV mutants containing varying numbers of single-nt mutations (12). One such mutant, designated Mut17, had four different single-nt mutations within the p28/p88 coding region. Among them, only one – C972U – did not alter the aa sequence of p28/p88. The remaining three mutations, C401U, G674A, and G2332U, altered identities of three aa residues (Fig. 1B). Specifically, alanine, arginine, and glycine residues at positions 113, 204, and 757 of p28/p88 were mutated to valine, histidine, and a premature stop codon, hence the names A113V, R204H, G757*, respectively (Fig. 1B).

Mut17 was originally identified in cells infected by a TCV replicon known as TCV-dMP_sg2R. This replicon harbored a deletion within p8/p9 movement protein (MP) genes, and an mCherry gene in place of the capsid protein gene (Fig. 1B). As a result, it replicated robustly in single cells, but could neither spread cell-to-cell nor assemble virus particles. When the four Mut17 mutations were re-introduced into TCV-dMP_sg2R together, the resulting mutant was unable to replicate in single cells, as judged by Northern blot hybridizations (Fig. 1C, lane 3). However, when each of the mutations was introduced into TCV-dMP_sg2R separately, the resulting A113V and N303N mutants replicated to levels similar to the mutation-free TCV-dMP_sg2R (Fig. 1C, compare lanes 4 and 6 with lane 2). By contrast, the R204H mutant replicated to very low, barely detectable levels (Fig. 1C, lane 5), whereas the G757* mutant RNA was undetectable (Fig. 1C, lane 7).

Intriguingly, while a faint sgRNA1 (sg1) band was detected in both TCV-dMP_sg2R and N303N samples (Fig. 1C, lanes 2 and 6), this band appeared to be missing in the A113V sample (Fig. 1C, lane 4). This observation prompted us to re-examine the impact of A113V mutation in wildtype (wt) TCV infections. To this end, A113V was introduced into an *Agrobacterium* (agro)-infiltration-compatible infectious clone of wt TCV and the resulting mutant was brought into *N. benthamiana* plants, with the wt TCV clone as control (Fig. 2). Moreover, to assess whether the inoculum dose had any effect on infection outcomes, we tested three different agro concentrations, with the respective optical densities (OD; at λ = 600 nm) adjusted to 1, 0.1, and 0.01, respectively. We were surprised to find that at 4 days post agro-infiltration (4 dpai), lower agro concentrations consistently correlated with higher viral gRNA levels in the agro-infiltrated local leaves (IL), for both wt TCV and the A113V mutant (Fig. 2A, compare lanes 2-4, 5-7). Therefore, lower inoculum dose probably facilitated more efficient replication in primary cells that received viral cDNAs.

**Figure 2.**
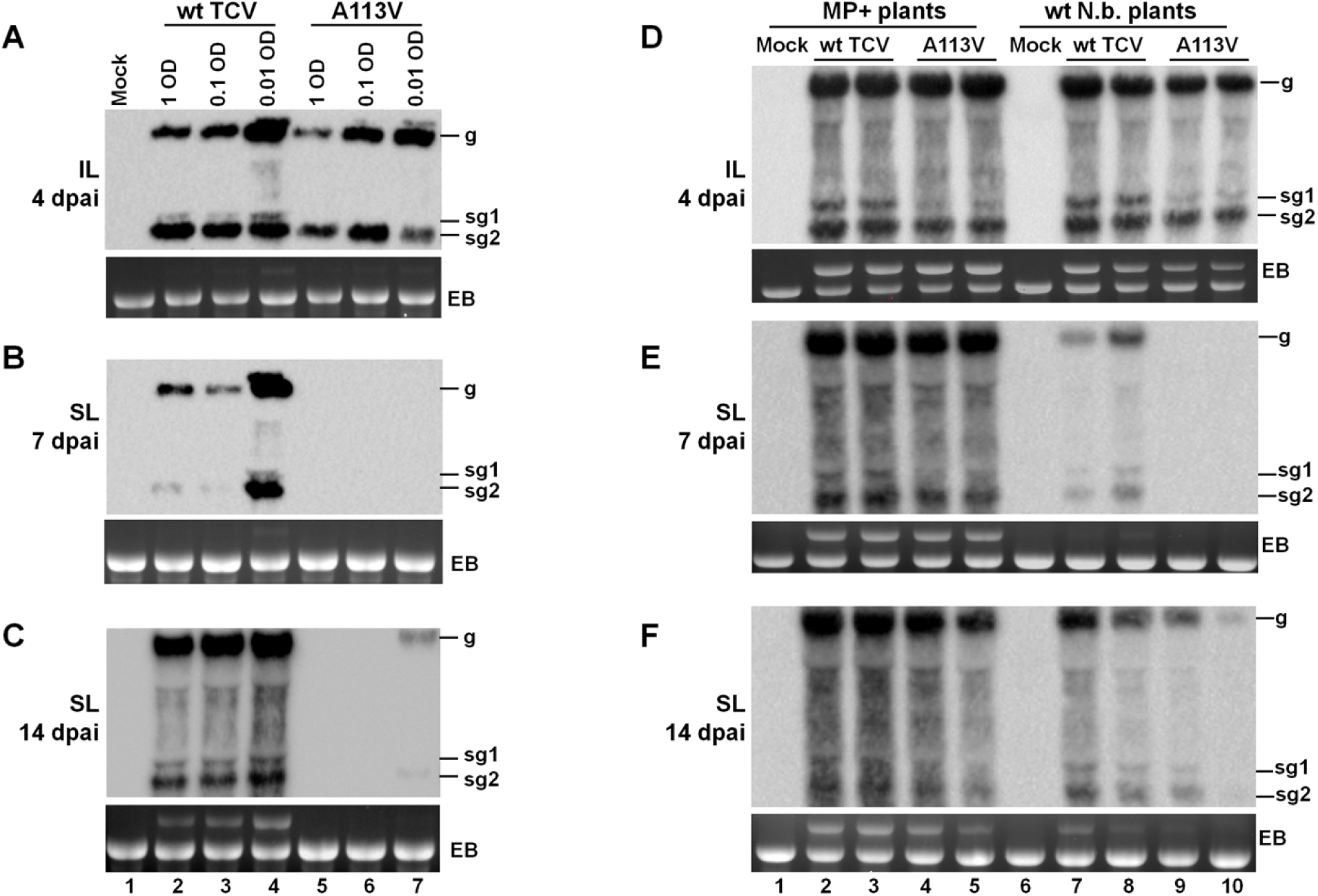
A113V diminishes sgRNA1 levels and delays TCV systemic spread. **A.** The A113V mutant failed to produce sgRNA1 in 4 dpai inoculated leaves (ILs) to levels detectable with NB. *Agrobacterium* suspensions harboring the A113V mutant, and the wt TCV control, were delivered into *N. benthamiana* leaves at three different concentrations (1, 0.1, and 0.01 OD). **B.** Viral RNA of A113V mutant was undetectable with NB in 7 dpai SLs. **C.** Viral RNA of A113V was marginally detectable with NB at 14 dpai (SLs) only in plants receiving the mutant at a very low dose (0.01 OD). **D - F.** MP+ plants expressing a heterogenous (RCNMV) MP mitigate the A113V defect in ILs and SLs. **D**. NB detection of wt TCV and A113V viral RNA levels in MP+ and wildtype *N. benthamiana* leaves at 4 dpai. Note the lower levels of sgRNA1 in lanes with the A113V mutant (lanes 4,5,9,10). **E and F.** wt TCV and A113V viral RNA levels in MP+ and wildtype *N. benthamiana* SLs at 7 dpai (E) and 14 dpai (F).

In contrast to wt TCV, the A113V mutant failed to produce detectable levels of sgRNA1 in ILs at all three inoculation doses (Fig. 2A, lanes 5-7). Consistent with the crucial role of sgRNA1 as mRNA for p8 and p9 MPs, viral RNAs of the A113V mutant were undetectable in systemic leaves (SLs) at 7 dpai (Fig. 2B, lanes 5-7). Furthermore, at 14 dpai, A113V RNAs were present only in SLs of plants inoculated with the 0.01 OD dose, and at low titers (Fig. 2C, lane 7; also see later). Consistent with the diminished viral RNA levels, the A113V-inoculated plants were symptom-free until at least 13 dpai, though some of them developed delayed symptoms afterwards (see later). Notably, the control plants inoculated with wt TCV developed clearly discernable symptoms by 7-8 dpai. Together, these results strongly suggest that while the A113V mutant replicated robustly in individual primary cells, its systemic spread was severely compromised, likely due to diminished sgRNA1 synthesis or stability, hence dearth of p8 and p9 MPs.

The delayed systemic spread of the A113V mutant could have been caused by other replication defects in addition to lower sgRNA1 levels. To assess this possibility, we tested the A113V mutant in transgenic *N. benthamiana* plants expressing MP of red clover necrotic mosaic virus (RCNMV), hereafter referred to as MP+ plants (13). The MP+ plants have been previously shown to complement TCV mutants with deletions in p8 and p9 (14). As shown in Fig. 2D, E, and F, in MP+ plants the A113V mutant yielded wt levels of viral gRNA in 4 dpai ILs as well as 7 and 14 dpai SLs. Of note, the sgRNA1 levels in A113V-infected tissues remained low or undetectable until at least 14 dpai. These results strongly suggested that reduction in sgRNA1 levels, and hence shortage of p8 and p9 MPs, were the primary culprit of systemic movement delay observed in A113V-infected wt *N. bethamiana* plants.

### A113V-infected plants exhibiting delayed symptoms contain viral variants harboring second-site mutations

By 21 dpai, some A113V-infected plants developed systemic symptoms of varying severity (Fig. 3A). We hence evaluated TCV RNA levels in SLs of these plants. For this analysis we selected four plants each from the 0.1 OD and 0.01 OD inoculum groups. As shown in Fig. 3B, viral RNA accumulation varied considerably in these plants, ranging from barely detectable in some plants (lanes 6-9), to wt levels in others (lanes 4, 5, 11). These results led us to speculate that the dominant TCV variants in some of the plants might have reverted to wt sequence or acquired compensatory second-site mutation(s).

**Figure 3.**
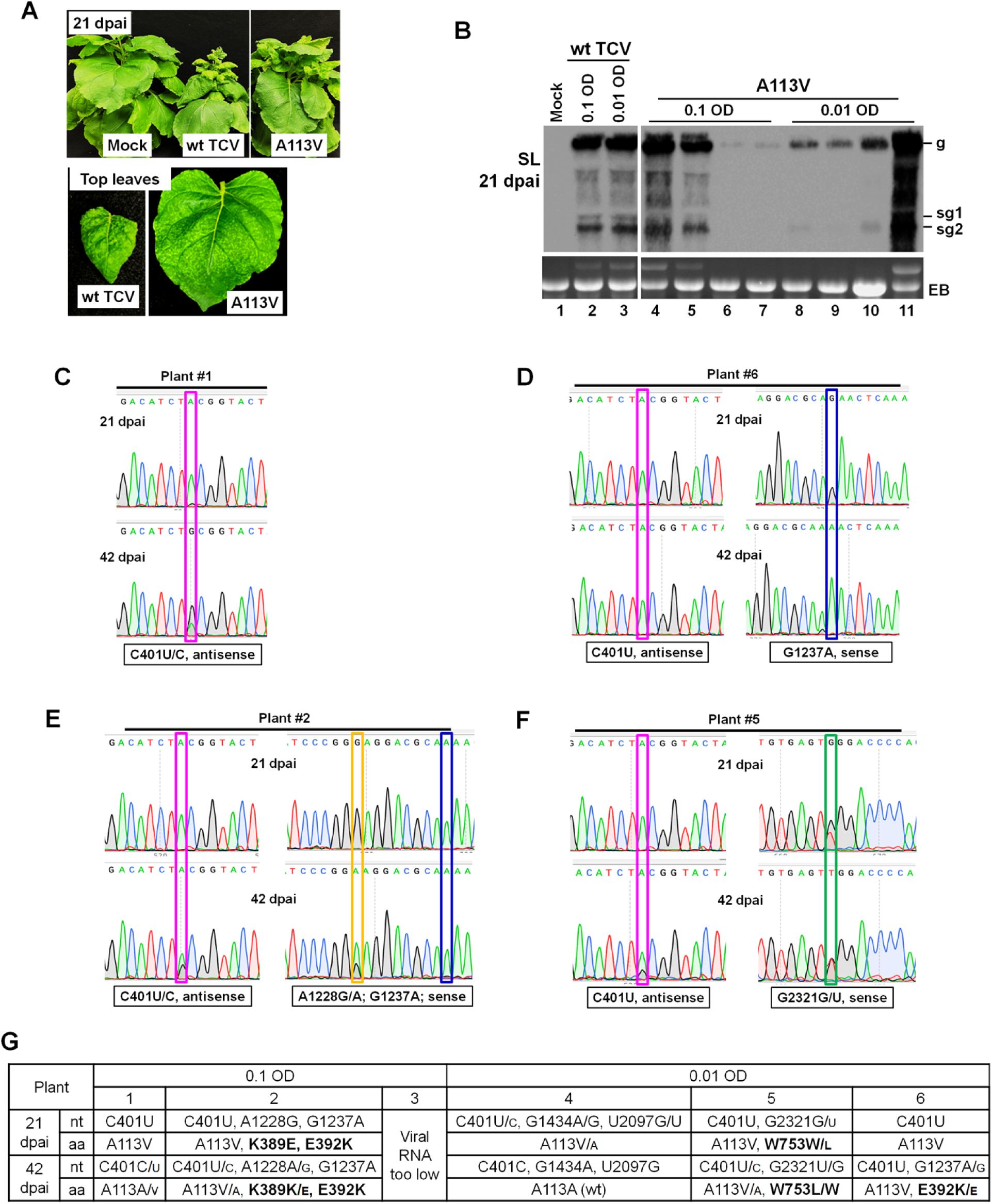
Identification of second-site mutations from SLs of A113V-infected *N. benthamiana* plants. **A.** Systemic symptoms of wt TCV and A113V-infected plants at 21 dpai. Bottom images reflect the actual size difference of two systemically infected leaves. **B.** NB detection of A113V viral RNA levels at 21 dpai, in 8 different plants. **C – F.** Chromatographs showing the emergence of reversion and/or second-site mutations in 4 different plants at 21 and 42 dpai. **G.** Summary of mutations identified from six different A113V-infected plants at 21 and 42 dpai.

To test this, we subjected the viral RNA in 3 plants of the 0.1-OD group and 3 plants of the 0.01-OD to reverse transcription (RT)-PCR to amplify the entire p28/p88 coding region. The amplified cDNA fragments were subsequently sequenced to identify dominant variants in the respective samples. The results showed that TCV cDNA of sufficient quantities was obtained in five of the six plants (except for plant #3; Fig. 3G). At 21 dpai, just two plants (#1 and 6) contained the original mutant with C401U (A113V) as the sole mutation (Fig. 3C and D; the U/C residues correspond to A/G peaks in antisense reads. Highlighted with purple boxes). However, the mutant was not stable in plant #1 because, by 42 dpai, the C401U mutation became a minor peak co-existing with the dominant C peak of wt TCV (Fig. 3C, bottom graph). On the other hand, although the C401U mutation remained stable in plant #6 by 42 dpai, an additional G1237A mutation emerged, altering aa position 392 from a glutamate to a lysine (E392K) (Fig. 3D; blue box).

Strikingly, the E392K mutation in plant #6 was also detected in plant #2, at the earlier 21 dpai time point (Fig. 3E). In this plant the E392K (G1237A) mutation was accompanied by another *de novo* mutation, K389E (A1228G; brown box), as well as the original A113V (C401U) mutation. More intriguingly, by 42 dpai, a minor C401 (wt) peak emerged under the dominant C401U peak (Fig. 3E). Meanwhile, a dominant A1228 (wt) peak overshadowed the A1228G (K389E) *de novo* mutation by 42 dpai (Fig. 3E). Nonetheless, the G1237A (E392K) *de novo* mutation was still stable (Fig. 3E). The fact that the E392K mutation emerged in two independent plants (#6 and 2. Fig. 3D and 3E) strongly suggested that this mutation may have compensated for the defect associated with the A113V mutation (more later).

The dominant variant in plant #5 was likewise noteworthy (Fig. 3F). At 21 dpai, a minor, yet unmistakable G2321U peak co-existed with the wildtype G2321. The G2321U mutation would have changed aa residue 753 from tryptophan to leucine (W753L). By 42 dpai, the G2321U peak rose to co-dominance with G2321 (wt). Curiously, by then the original C401U (A113V) mutation was also beginning to be accompanied by a minor C401 (wt) peak. Thus, W753L was probably another mutation that compensated for the A113V defect. In summary, two new mutations, E392K and W753L, may have been selected for their ability to rescue the loss of sgRNA1 in the A113V mutant infections.

Finally, the dominant variant in plant #4 incurred two second-site mutations, G1434A and U2097G, in addition to a minor C401 (wt) peak under the dominant C401U peak, at 21 dpai (Fig. 3G and data not shown). However, neither of them altered any aa residue. Moreover, in this plant the C401U mutation reverted to wt by 42 dpai. Together these data suggested that the sgRNA1 loss caused by the A113V mutation required additional compensatory mutations, or simple reversion to wt, in order to restore systemic spread to the virus. It is further noteworthy that these compensatory mutations occurred within p88 RdRp, at positions distant from the original A113V mutation at the primary aa sequence level.

### E392K and two double mutants yield sgRNA1-like alternative sgRNAs and alleviate A113V-caused delays in TCV systemic spread

To resolve whether the second-site mutations could indeed rescue the A113V defects, we introduced these mutations, including K389E, E392K, and W753L, as well as K389E-E392K double mutations (designated KEEK for simplicity), into wt TCV and A113V backbones (Fig. 4A). Northern blot hybridization results revealed that in 4 dpai ILs, these mutants produced drastically different levels of viral gRNA and sgRNAs, and novel sgRNA species in some cases. It should be noted that in this set of experiments the A113V mutant did accumulate low levels of sgRNA1 (Fig. 4B, compare lanes 13-14 with 3-4). Consistent with earlier observations, A113V systemic movement was delayed as its RNAs were undetectable in 7 dpai SLs, and detectable in just one of the two plants examined at 14 dpai (Fig. 4C and D, lanes 13-14). More tellingly, sequence analysis revealed that in this plant the A113V mutation has reverted to wt TCV (Fig. 4E, sample #14).

**Figure 4.**
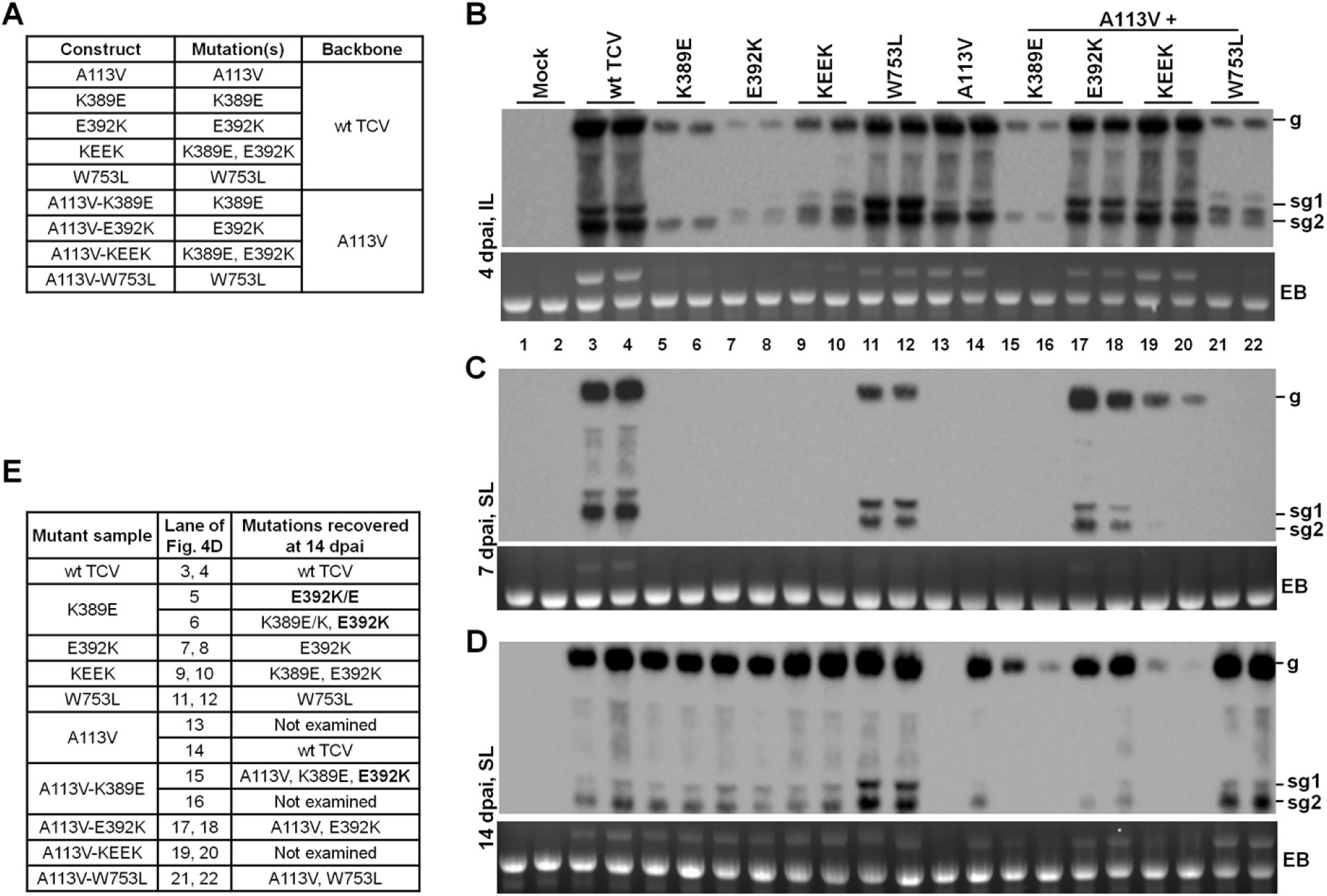
*In planta* fate of second-site mutations by themselves, and in combination with A113V. **A.** A list of 8 mutants created. **B – D.** NB determination of viral RNA levels for the 8 mutants in ILs at 4 dpai (B), in SLs at 7 (C) and 14 (D) dpai. **E.** Dynamic changes of the mutations in 14 dpai descendants of the 8 mutants as determined by sequencing of RT-PCR products.

The K389E mutation by itself completely abolished sgRNA1, but also substantially decreased the levels of gRNA and sgRNA2 in 4 dpai ILs (Fig. 4B, compare lanes 5-6 with 3-4). Combining K389E with A113V failed to resurrect sgRNA1, and instead decreased gRNA and sgRNA2 levels further (Fig. 4B, lanes 15-16). Moreover, neither K389E single mutant nor K389E-A113V double mutant could be detected in SLs at 7 dpai, reflecting delayed systemic spread (Fig. 4C, lanes 5-6, 15-16). We hasten to note that the K389E mutation was never isolated as a stand-alone mutation in earlier experiments. Rather, it was detected in one of the A113V-infected plants along with E392K and A113V (Fig. 3). Interestingly, the K389E single mutant, but not the A113V-K389E double mutant, accumulated high levels of TCV RNA in 14 dpai SLs (Fig. 4D, lanes 5-6, 15-16).

Sequence analysis of RT-PCR products revealed that by 14 dpai, in one of the K389E-infected plants the K389E mutation reverted to wt, and a new E392K mutation simultaneously emerged (Fig. 4E, sample #5). In the second plant, the partially reverted K389E was accompanied by the same E392K mutation (Fig. 4E, sample #6). Finally, the A113V-K389E double mutant replicated poorly in 14 dpai SLs and, in the sample subjected to sequence analysis (Fig. 4E, sample #15), it likewise acquired the E392K mutation. Therefore, the K389E mutation by itself was likely severely deleterious. Conversely, the E392K mutation was likely enriched due to its ability to compensate for the defects of K389E, as well as that of A113V.

The E392K single mutant was even more deleterious than K389E (Fig. 4B, compare lanes 7-8 with 5-6) in 4 dpai ILs, as mutant gRNA was barely detectable, and its sgRNA2 levels were lower still (Fig. 4B, lanes 7 and 8). More strikingly, sgRNA1 was replaced by two new sgRNA species, migrating slightly slower and faster than wt sgRNA1, respectively (Fig. 4B, lanes 7-8). For convenience, hereafter we refer to the slower-migrating species as sgRNA1a, the faster-migrating one sgRNA1b, characterization of which to be described later. Notably, the sgRNA1b band intensity was higher than sgRNA2, reflecting a perturbation of sgRNA relative abundances. Both sgRNA1a and 1b were also evident in ILs receiving KEEK and A113V-W753L mutants at 4 dpai (Fig. 4B, lanes 9-10; 21-22), and they were consistently observed in at least three independent repeat experiments.

Contrary to K389E, the E392K defect was dramatically corrected in the A113V-E392K double mutant (Fig. 4B, compare lanes 17-18 with 15-16). Specifically, viral RNAs of A113V-E392K double mutant accumulated to substantially higher levels than the E392K single mutant, and the sgRNA1a and 1b bands also disappeared (Fig. 4B, compare lanes 7-8 and 17-18). Instead, the wt sgRNA1 band reappeared at a relative abundance similar to wt TCV. Accordingly, the A113V-E392K double mutant, but neither the A113V nor E392K single mutant, accumulated viral RNA levels in 7 dpai SLs to near wildtype levels (Fig. 4C, compare lanes 17-18 with 7-8 and 13-14). Of interest, the E392K single mutant, though delayed, was able to reach 14 dpai SLs at relatively high levels, with the original mutation stably retained (Fig. 4D, lanes 7-8; Fig. 4E). Likewise, the A113V-E392K mutant also retained both mutations for at least 14 days (Fig. 4D, lanes 17-18; Fig. 4E). In short, the A113V-caused defects, both the loss of sgRNA1 and delay in systemic spread, were successfully rescued by the E392K mutation.

The KEEK double mutant accumulated viral RNAs to levels higher than K389E and E392K alone, but still lower than wt TCV, in 4 dpai ILs (Fig. 4B, lanes 3-10). Intriguingly, KEEK retained the sgRNA pattern of E392K (sgRNA1a, 1b, sgRNA2) but not K389E (sgRNA2 only). Thus, it appears likely that K389E was primarily enriched in the E392K context to compensate for the E392K-caused replication inefficiency. Consistent with this interpretation, in 4 dpai ILs the A113V-KEEK triple mutant accumulated more gRNA and sgRNA2 than the A113V-E392K double mutant but modestly less sgRNA1 (Fig. 4B, compare lanes 19-20 with 17-18). Perhaps due to the lower sgRNA1 levels, the A113V-KEEK triple mutant was less efficient at systemic spread, and gradually diminished in SLs (Fig. 4C and D, lanes 19-20).

The W753L single mutant accumulated the same RNA pattern – gRNA, sgRNA1, and sgRNA2 – as the wt TCV in 4 dpai ILs. However, the sgRNA1 levels rose substantially to levels exceeding that of sgRNA2 (Fig. 4B, lanes 11-12). Among the four single mutants (A113V, K389E, E392K, W753L), W753L was the only one detectable in 7 dpai SLs, albeit at lower levels than wt TCV. Notably, the relative over-abundance of sgRNA1 of W753L persisted in SLs until at least 14 dpai (Fig. 4C and D, lanes 11-12). On the other hand, the A113V-W753L double mutant, contrary to A113V-E392K and A113V-KEEK, replicated much worse than A113V or W753L single mutant (Fig. 4B, compare lanes 21-22 with 11-14). Curiously, the A113V-W753L also accumulated sgRNA1a and 1b, with their sizes and relative ratio resembling that of E392K and KEEK mutants (Fig. 4B, compare lanes 21-22 with 7-10). Thus, loss of wt sgRNA1 and concurrent emergence of sgRNA1a and 1b appeared to be a shared response to certain types of structural disturbance, rather than a specific response to particular aa mutations.

Conversely, the W753L mutation in the A113V background was probably a temporary compensation for the loss of sgRNA1. This is because, unlike W753L single mutant, the A113V-W753L double mutant still exhibited delayed systemic infection (Fig. 4C, lanes 21-22). Nevertheless, this mutant accumulated to wt levels in 14 dpai SLs (Fig. 4D, lanes 21-22), and retained both mutations (Fig. 4E, samples #21 & 22). In summary, W753L appears to compensate for the A113V defects by increasing the levels of sgRNA1 only.

### sgRNA1a and sgRNA1b have different 5’ proximal sections, but nearly identical remaining sequences

We next wondered how various second-site mutations compensated for reduced sgRNA1 levels and alleviated the delay in viral systemic spread caused by the A113V mutation. Since three different sets of mutations – E392K, KEEK, and A113V-W753L – led to the emergence of two alternative sgRNA species – sgRNA1a and sgRNA1b – in lieu of wt sgRNA1, we sought to uncover possible compensation mechanisms by resolving the identities of these novel sgRNAs. We hasten to note that these alternative sgRNA species were detectable in ILs only, despite the persistence of some of the mutations in 14 dpai systemic leaves (Fig. 4D, lanes 7-8, 9-10, 21-22; Fig. 4E). Nevertheless, their identities should still be informative as they probably represented immediate, though not necessarily lasting, natural selection response to the original A113V mutation.

Previous studies suggested that replicating TCV gRNA and sgRNAs frequently incurred covalent linkages between the 5’ and 3’ termini of the same or two different copies of the same RNA, forming circularized or dimeric intermediates (15); (Khemsom and Qu, unpublished). We took advantage of these end-joined forms of sgRNAs for simultaneous mapping of the 5’ and 3’ ends of alternative sgRNAs, using a pair of divergent RT-PCR primers (TCV-3872F and -2741R. Fig. 5C, orange arrowheads). Fig. 5A shows the RT-PCR products programmed with RNA samples extracted from ILs of wt TCV and two double mutants – KEEK and A113V-W753L. The 3872F and 2741R primer pair should generate RT-PCR products only when the sgRNA templates were in circular or dimeric forms, yielding a 594-bp fragment from sgRNA1 (Fig. 5B & C; the 183-bp dull blue fragment at the 3’ end fused to the 411-bp red fragment corresponding to the 5’ end of sgRNA1), and a 319-bp fragment for sgRNA2 (Fig. 5B & C, the 183-bp dull blue fragment fused to the136-bp pink fragment).

**Figure 5.**
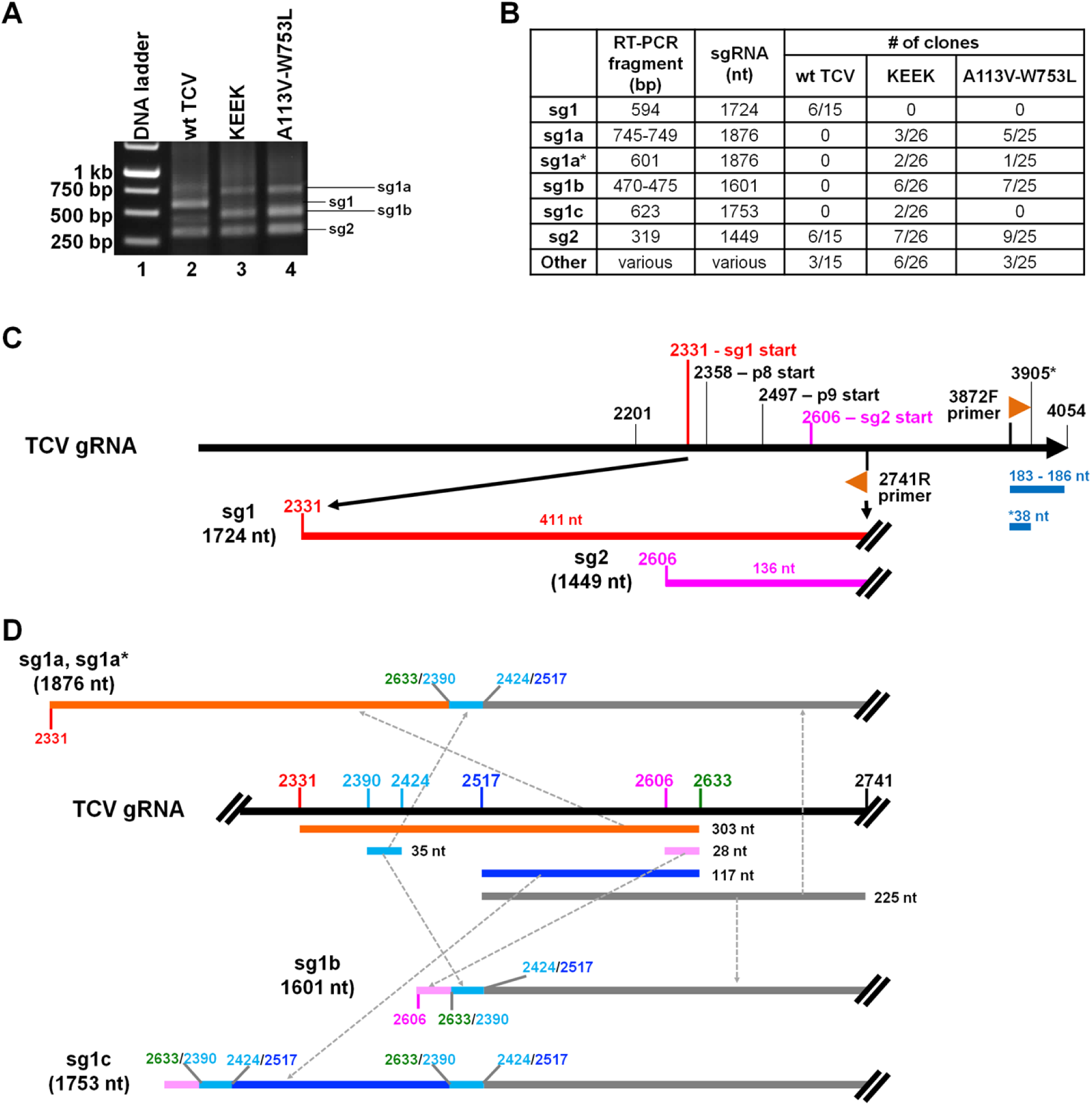
Molecular characterization of alternative sgRNA1 (sg1) species. **A.** RT-PCR products generated with a pair of primers that enrich circular or dimeric forms of sgRNAs, from three representative virus-infected plant RNA samples, infected with wt TCV, KEEV, and A113V-W753L, respectively. The tissues were SLs collected at 14 dpai. **B.** Summary of sequencing results, showing various unique clones and their respective numbers. See C and D for the composition of three representative classes of clones. **C.** Schematic depiction of expected RT-PCR fragments derived from wildtype sgRNA1 and 2 (sg1 and sg2). **D.** Assembly of sgRNA1a, 1a*, 1b, and 1c based on sequencing results.

Notably, the KEEK and A113V-W753L samples both yielded 3 products of similar sizes – approximately 750, 500, and 300 bp. Note the sizes of these RT-PCR products were consistent with them being derived from sgRNA1a, 1b, and sgRNA2 (Fig. 4), with the 300-bp product migrating at the position of sgRNA2-specific product in the wt TCV lane (Fig. 5A, lane 2-4; Fig. 5C). We then cloned these products into a plasmid vector and sequenced multiple clones to resolve the identities of these fragments. We were surprised to find that the similar-sized products derived from KEEK and A113V-W753L samples shared nearly identical sequences.

Several classes of clones have been recovered. The first class consisted of 8 clones – 3 from the KEEK sample and 5 from the A113V-W753L sample, harboring nearly identical inserts of 745-749 bp (Fig. 5B). Four of the 8 inserts were 747 bp, and the minor size variations in remaining 4 clones were due to differences at the 3’-5’ junctions. Excluding the183-186 bp originating from the TCV 3’ terminus (position #3872-4054 plus a few extra C or Gs), the remaining 563 bp was made up of three partially repetitive, discontinuous TCV sections (Fig. 5D, top diagram): a 303-bp 5’ proximal section (Fig. 5D, orange line) that began at the authentic sgRNA1 start site (position 2331) and ended at position 2633 of TCV gRNA; followed by a 35-bp section spanning positions 2390-2424, thus internal to the first section (Fig. 5D, light blue lines); as well as a 225-bp 3’ proximal section spanning positions 2517-2741 (Fig. 5D, gray lines). Assuming no additional changes downstream of position 2741, these chimeric inserts would have been derived from an sgRNA1a of 1,876 nt, surpassing wt sgRNA1 (1,724 nt) by 152 nt.

The second class consisted of 3 clones – 2 from the KEEK sample and 1 from the A113V-W753L sample – had inserts with a shorter, 38-bp 3’ terminal section (positions 3872-3909), but the same 5’ 563 bp as the first class of clones. The corresponding alternative sgRNA would be of same size as sgRNA1a, hence was named as sgRNA1a* (Fig. 5B and C). Collectively 5 of the 26 KEEK clones, and 6 of 25 A113V-W753L clones, arose from the nearly identical, 1,876-nt sgRNA1a. Finally, it is worth reiterating that the 5’ termini of sgRNA1a and 1a* were identical to that of wildtype sgRNA1.

The third class of clones were recovered 13 times – 6 from KEEK and 7 from A113V-W753L samples Fig. 5B; Fig. 2D, sg1b diagram). Their inserts ranged from 470-475 bp, with the minor size differences again mapping to the 3’-5’ junctions. Excluding the 183-to-188-bp section originating from TCV 3’ end, the remaining 287 bp also comprised three discontinuous sections. The 28-bp 5’ proximal section (Fig. 5D, light purple line) initiated at the authentic sgRNA2 start site (position 2606) and, intriguingly, ended at the same 2633 position as sgRNA1a and sgRNA1a* (Fig. 5D). Moreover, the next 35-bp section (Fig. 5D, light blue line) and the last 225-bp section (Fig. 5D, grey line) were likewise shared by sgRNA1a/1a*-derived cDNA. These inserts would predict an alternative sgRNA1b of 1,601 nt, 123-nt shorter than wt sgRNA1, but 152-nt longer than wt sgRNA2 (Fig. 5D).

The fourth class of clones were recovered twice, from the KEEK sample only. Their insert of 623-bp was particularly interesting (Fig. 5D, sg1c diagram). Excluding the 183-bp section derived from TCV 3’ terminus, the remaining 440 bp represented reiterated RNA recombination events at the same hotspots. It contained the same two 5’ sections as sgRNA1b. But the next, 117-bp section represented yet another pause at position 2633, and a second recruitment of the 35-bp spanning positions 2390-2424. These additional recombination events gave rise to the alternative sgRNA1c that was 1,753 nt in size, 29-nt longer than wt sgRNA1. Taken together, these results resolved the identities of sgRNA1a and 1b that emerged in cells infected by two distinct double mutants – KEEK and A113V-W753L.

The deduced sequence of sgRNA1a could readily template the translation of wt p8 (Suppl Fig. 1A). Furthermore, despite the 35-nt insertion (light blue lines in Fig. 5D) and partial duplication of the p9 coding sequence, it could support undisrupted translation that began with an AUG internal to original p9 coding sequence (but in a different reading frame. Suppl Fig. 1A), read through the 35-nt insertion, and then continue into the N-terminally truncated p9 coding sequence downstream. The resulting p9 variant replaced the N-terminal 7 aa of wt p9 with a 29-aa novel N-terminus. Therefore, sgRNA1a could potentially rescue the loss of sgRNA1 by templating the translation of a wt p8 and an N-terminally altered p9. By contrast, sgRNA1b (and sgRNA1c) was unlikely to have rescued the loss of sgRNA1 (Suppl Fig 1B). This is because it lacked most of the p8 coding sequence. Even though a putative coding frame for an N-terminally truncated and mutated p9 variant could be identified (Suppl Fig 1B), that frame did not contain any AUG start codon, hence was unlikely to translate this p9 variant.

### Amino acid changes in compensatory mutants do not cluster with A113 in predicted three-dimensional (3-D) structure of p88

To elucidate how A113V, K389E, E392K, and W753L mutations influence TCV replication and sgRNA transcription, we modeled the 3-D structure of a putative TCV replication complex comprising p28, p88, and a 150-nt single-stranded RNA (ssRNA) template annealed to a 26-nt RNA product. To maintain consistency with the symmetry observed in the flock house virus (FHV) and Chikungunya virus replication factories (16), the modeled complex included one p88 and eleven p28 molecules, reflecting the higher abundance of p28 during infection. AlphaFold3 predicted a 12-member conical ring structure, with the p88 RdRp domain positioned centrally within the channel (Fig. 6A, B, C). The N-terminal 20 aa of both p28 and p88 were predicted to form an amphipathic α-helix (i.e. α1) located at the broader end of the conical ring, oriented with the hydrophobic face outward (Fig. 6B). This orientation suggests that the broader end of the ring likely anchors to the host membrane to facilitate the assembly of replication vesicles. The modeled RNA molecule, representing gRNA with both single-stranded and double-stranded regions, was predicted to wrap along outer surface of the conical ring before threading into the RdRp active site at the central channel. Notably, similarly shaped conical ring structures were predicted regardless of the numbers of p28 (i.e. 11 to 18) and p88 (i.e. 1 to 2) subunits specified in the AlphaFold3 input, with or without RNA.

**Figure 6.**
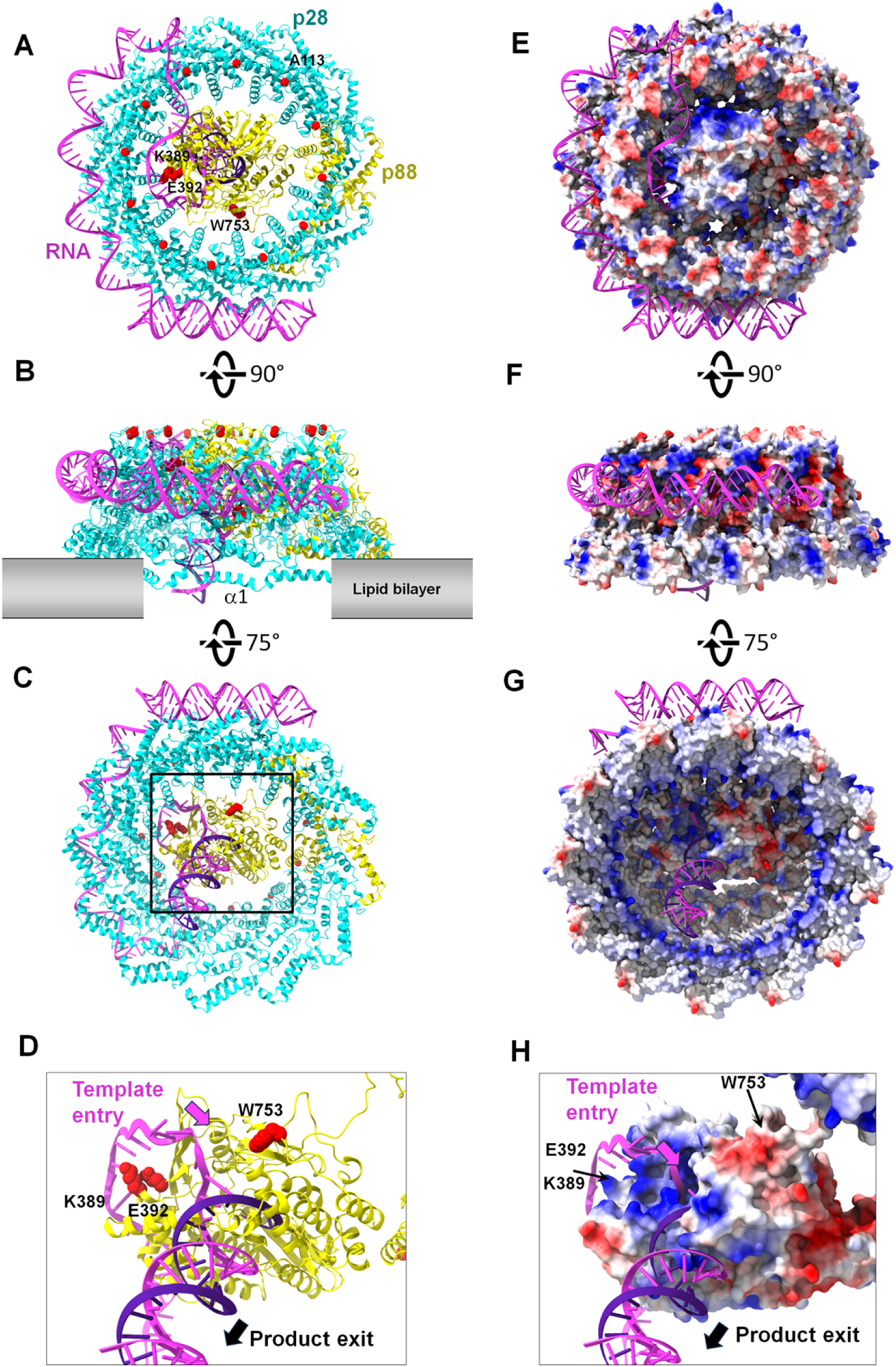
Molecular modeling of the TCV replication complex. **A – C**. Structural model of the TCV replication complex. The model contains eleven p28 (cyan), one p88 (yellow), and a 150-nt RNA template (magenta) annealed to a 26-nt RNA product (purple). Residues A113, K389, E392, and W753 are highlighted in red spheres. Three orientations - top, side, and bottom - are shown to illustrate the overall organization of the complex. The first α-helix (α1) of both p28 and p88 is amphipathic and is likely responsible for membrane insertion of the replication assembly. **D**. The RdRp active site view. The p88 RdRp domain is shown with a template-product duplex derived from Alphafold3-based structural modeling. Panel D provides a magnified view of the boxed region in panel C. **E – H**. Electrostatic surface of the replication complex and the RdRp domain. Molecular surface representations are colored by electrostatic potential with positive in blue and negative in red.

Inspection of the conical ring revealed that the residue A113 is positioned within a protruding loop situated at the narrower end of the ring (Fig. 6A, B, C). AlphaFold3 modeling predicts that this residue contacts the RNA template (e.g. gRNA) during synthesis of the (-) RNA progeny (Fig. 6A), potentially facilitated by the regularly spaced, positively charged surface patches distributed along the outer periphery of the complex (Fig. 6E, F, G). Alternatively, the loop containing A113 may serve as a docking site for a host factor that mediates premature termination of RNA replication to generate (-) sgRNA1 molecules. The A113V substitution could alter either the template RNA binding or host factor interaction in such a way that selectively impairs synthesis of (-) sgRNA1 but not (-) sgRNA2.

Interestingly, the three residues affected by second-site mutations, K389, E392, and W753, are located within the p88 RdRp domain, spatially distant from A113. To explore how mutations at these positions influence sgRNA synthesis, we further modeled the template:product duplex from the HIV reverse transcriptase ternary complex into the p88 RdRp domain (Fig. 6D, H). The resulting model shows that K389 and E392 lie near the outer rim of the template entry channel, where they directly interact with both the phosphate backbone and nucleotide bases of the RNA template. It is therefore plausible that the K389E and E392K substitutions altered local electrostatic or hydrogen-bonding interactions in a sequence-dependent manner, leading to transient stalling of RdRp and facilitating template switching at sequence-specific hotspots during (-) sgRNA synthesis. W753, located on the opposite side of the template entry channel, may likewise interact with the template RNA to modulate pausing and template switching to produce sgRNA1a, and 1b.

## Discussion

The biogenesis of sgRNAs of (+) RNA viruses is known to be dependent on RNA secondary structures in the vicinity of sgRNA initiation sites, as well as specific sequence motifs located at the 3’ termini of sgRNA (-) strands (10, 17–19). A large group of viruses, including all viruses in the family *Tombusviridae* to which TCV belongs, use a premature termination mechanism that enlists secondary structures formed by viral RNA genomes to terminate (-) strand synthesis at specific sites, thereby releasing (-) strands of defined lengths, which can then be used as templates for (+) strand sgRNA synthesis (7, 20, 21). These released (-) strands mostly also contain conserved sequence motifs at the 3’ ends, known as promoters, that recruit RdRp for (+) strand polymerization (22, 23). However, whether viral RPs contribute to the determination of number and sizes of sgRNAs is not well understood. Among the few reported studies, Wu and White (20) showed that the identities of five C-terminal aa residues of p92, the RdRp of tomato bushy stunt virus (TBSV), were critically important for the accumulation of sgRNA2 of that virus. However, since those authors used a TBSV mutant that no longer produced sgRNA1, it is not known whether these five aa regulated the synthesis (transcription) of one or both sgRNAs. A follow-up study by Gunawardene and colleagues (21) identified several single aa changes of TBSV p92 that either abolished viral replication, or transcription of sgRNAs only. These single aa changes were introduced into a C-proximal region of p92 conserved among viruses of *Tombusviridae*. It was found that 70% of mutants located inside a 19-aa domain preferentially affected sgRNA2 production. However, these mutants were only interrogated in single protoplast cells. As a result, it is unknown if these defects could be rescued by compensatory changes elsewhere in p92. Separately, a more recent study by Martin and colleagues (24) found that certain mutations in the Chikungunya virus RdRp compromised the accumulation of capsid protein and virions, suggesting, albeit indirectly, that these mutations might compromise the synthesis of the sgRNA templating capsid protein translation.

The current study investigated four naturally occurring mutations of TCV p28/p88 that, alone or combined, perturbed the biogenesis of TCV sgRNA1. Specifically, A113V and K389E reduced sgRNA1 to low or undetectable levels, whereas W753L boosted sgRNA1 to levels higher than sgRNA2, thus upsetting the relative abundance of the two. Furthermore, E392K, KEEK, and A113V-W753L engendered a novel three-sgRNA pattern by decreasing the levels of sgRNA2, and replacing wt sgRNA1 with two alternative sgRNAs – sgRNA1a and 1b. Notably, while the E392K single mutant accumulated poorly in infected cells, this defect was readily corrected in the A113V-E392K double mutant, indicating that these two mutations are mutually compensatory. Collectively these findings suggest that some of the mutations (A113V and K389E) may have disrupted the synthesis of the (-)-strand intermediates of wt sgRNA1, whereas other mutations or mutation combinations (E392K, KEEK, and A113V-W753L) may have selectively favored the production of (-)-stranded recombination products (sgRNA1a and 1b) that nevertheless shared the same terminal features with wt sgRNA1 or sgRNA2. Below we discuss how such preferential synthesis of different (-) strands could have been realized.

### Recombination events giving rise to alternative sgRNA1a and 1b likely have occurred during (-)-strand synthesis

We showed in Fig. 5 that sgRNA1a, 1a*, and 1b shared the same two recombination junctions that fused three discontinuous sections of TCV genome. Their size difference can be straightforwardly attributed to their different 5’ termini: sgRNA1a and 1a* possessed the 5’ terminus of wt sgRNA1, whereas sgRNA1b possessed that of wt sgRNA2. These characteristics are in turn best explained if recombination took place during (-)-strand synthesis (25–27). As schematically depicted in Fig. 7, mutations like E392K, KEEK, and A113V-W753L likely caused (-) strand synthesis to pause more frequently at certain sites of TCV genome. One such site is position 2517, where strong base-pairing between GGGGG in the template and the newly copied CCCCC may stall RdRp (Fig. 7, Step 1). The fragment released, possibly after adding an extra, non-templated C, would then re-attach to position 2424 where its 3’-terminal CCCCCC could anneal to GGCGG with one mismatch (Fig. 7, Step 2). After a short, 35-nt extension, the (-) strands apparently encountered a second site of frequent pauses, releasing a product with a 12-nt terminal motif (5’-AGAGAGUUGUAG-3’) (Fig. 7, Step 3). Incidentally, this 12-nt motif was 100% complementary to another template location near position 2633, permitting the released (-) strands to re-anneal at this downstream position, followed by continued copying to the authentic termination signals of wt sgRNA1 and 2, thereby completing the synthesis of (-)-strand intermediates of sgRNA1a and 1b, respectively (Fig. 7, Step 4).

**Figure 7.**
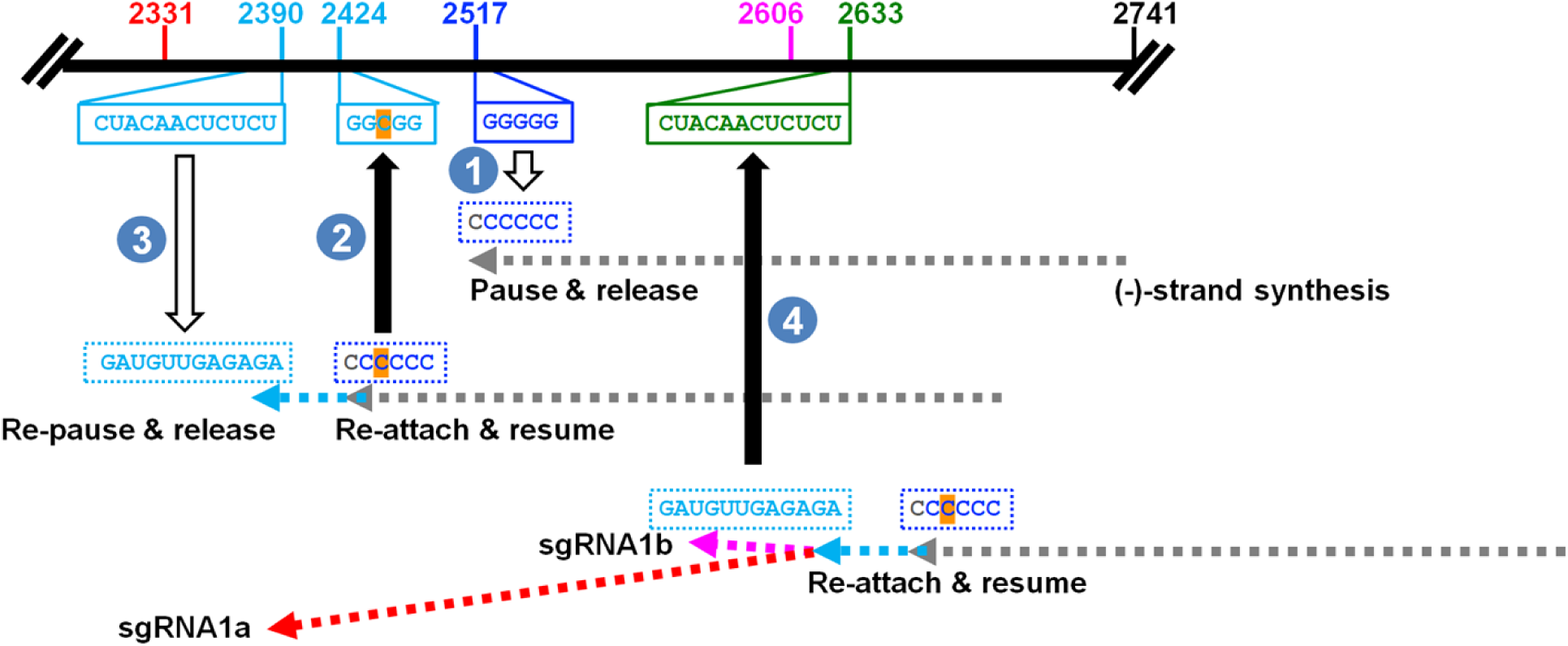
A model for the genesis of alternative sgRNA1 species.

### How might the A113V, K389E, E392K, W753L mutations, or combinations of them, alter the pattern of (-) sgRNA synthesis?

Replication of TCV gRNA, as well as transcription of its sgRNAs, are known to take place inside membranous spherules formed by invagination of the mitochondrial outer membrane (28, 29). These replication compartments morphologically resemble those described for brome mosaic virus (BMV) and FHV, which have been more thoroughly studied (30–35). Such spherules are typically induced by virus-encoded RPs, such as p28 and p88 in TCV, 1a and 2a in BMV, and Protein A in FHV. Recent cryo-EM studies of FHV replication complexes revealed a crown-like structure in which 24 copies of protein A organize into two stacks of 12-mer rings of, forming the spherule neck channel (32, 33).

The putative TCV replication complex we obtained through AlphaFold3 modeling consists of eleven copies of p28 and a single copy of p88, with the p88 RdRp domain positioned within the central channel of the complex (Fig. 6). This assembly is likely situated at the neck of the spherule invagination on the mitochondrial outer membrane. Interestingly, our model suggests that the first amphipathic α-helix of p28/p88 likely anchors the complex to the membrane. This is consistent with our earlier finding that deletion of the N-terminal 36 aa of p88 led to dramatic over-accumulation of a very short, 283-nt TCV sgRNA known as tiny TCV sgRNA or ttsgR - indicative of defective or malformed spherule structures (15). While the exact molecular stoichiometry of p28 and p88 within actual TCV replication complexes remains unresolved, previous work from our laboratory indicates that maintaining an appropriate p28:p88 ratio is critical. Zhang and colleagues (15) demonstrated that p88 overexpression caused a TCV replicon to generate markedly lower levels of gRNA, but elevated levels of both sgRNA1 and sgRNA2. Similarly, the relative abundance of BMV-encoded 1a and 2a RPs has also been shown to be essential for BMV replication (30, 36).

Within the predicted replication complex, A113, K389, E392, and W753 are all solvent-exposed and do not appear to participate in interactions among different subunits. As a result, these mutations are unlikely to have altered the p28:p88 ratios or disrupted the assembly of the replication complex. Instead, the increased abundance of sgRNA1a and 1b in these mutants likely resulted from preferred synthesis. Based on the positions of these residues in the modeled replication complex (Fig. 6), we speculate that A113V, being located at the narrower end of the conical ring, may alter the interaction between the replication complex and either the template RNA or potential host cofactors, thereby interfering the premature termination of (-) RNA synthesis that is required to generate (-) sgRNA1. In contrast, K389, E392 and W753, three residues found to be mutated *de novo* in A113V-infected tissues, lie near the template entry channel in the p88 RdRp domain. As a result, mutations at these sites could affect template engagement in a way that promotes stalling at sequence hotspots. Such stalling may facilitate template switching, giving rise to alternative (-) sgRNA1a, 1b, or 1c. Notably, premature termination of TCV (-) sgRNAs also relies on stalling conferred by RNA secondary structures, at specific positions (17).

In addition, compensation conferred by the W753L mutation might not be solely due to the W-to-L aa change – the underlying G2321U nucleotide change might have contributed as well. As noted earlier, viruses like TCV synthesize (-) sgRNA intermediates via a premature termination mechanism that relies on RNA secondary structures in the (+)-stranded templates to stall (-) strand elongation, thereby defining the 3’ termini of these (-) strands. It is thus reasonable to assume that stronger RNA secondary structures cause more frequent stalling of (-) strands, leading to more (-) strands that template more (+) strands. Incidentally, the single G2321U nt mutation underlying W753L also converted a G-G mismatch to a G:U base pair in the RNA secondary structure responsible for generating sgRNA1 (-) strands (Suppl Fig. 2) (37), causing modest stabilization of this structure. Such structural stabilization could conceivably lead to more sgRNA1 (-) strands.

### Why were sgRNA1a and 1b absent in systemic leaves of plants infected with E392K, KEEK, and A113V-W753L?

Interestingly, the three-sgRNA pattern observed in ILs of E392K, KEEK, and A113V-W753L infections was not detected in SLs of the same plants, despite the apparent stability of some of the mutations (Fig. 4E). However, it should be noted that systemic spread of E392K, KEEK, and A113V-W753L mutants were still delayed, with their RNAs undetectable in SLs until 14 dpai. We speculate that the different physiological states of cells (e.g. availability of host factors or different posttranslational modifications, etc) in ILs and SLs might permit mutant viruses to synthesize sgRNA1 and sgRNA2 more readily in SLs notwithstanding the mutated RPs. More specifically, cells in ILs are mostly mature source cells exporting energy-rich carbohydrates, whereas those in SLs are immature sink cells receiving carbohydrates. Additionally, there might have been other mutations outside the RP coding region of TCV genome that could have restored the formation of wt sgRNA1-accomodating replication compartments.

Finally, we wish to reiterate that the rescue afforded by compensatory mutations was by no means perfect. Rather, they provide transient relief for the A113V defects. This then allows their descendants to replicate to higher copy numbers in the subsequently invaded cells than sister lineages not containing the compensatory mutations. Such temporary solutions need not to be perfect – they simply need to be comparatively more fit then and there, permitting emergence of revertant or additional new mutants with superior compensatory capacities. On the other hand, embarking on the trajectory of harboring both original (A113V) and compensatory (e.g. E392K) mutations also means that the new mutant is now more difficult to revert, thus possibly trapping it in a suboptimal state for some lengths of time. To summarize, our current study identified several naturally occurring mutations that specifically affect one of the viral sgRNAs, and examined the natural selection remedies that compensated for their defects. These findings are expected to inform strategies that target viral sgRNA synthesis for controlling viral diseases of plants, animals, and humans.

## Material and Methods

Detailed descriptions of all materials and methods, including plasmid constructs used, plants and plant care, the virus inoculation procedure (agro-infiltration), viral RNA detection via Northern blot hybridization, RT-PCR and sequencing, and structural modeling of the TCV replication complex, are provided in the Supporting Information. Sequences of all primers are available upon request.

### Data sharing plan

All data collected throughout the current study have been provided in the manuscript text and Supplemental Information (SI) Appendix. Readers are encouraged to contact us with any questions.

## Acknowledgements

We thank the USDA ARS Maize and Soybean Viruses Group and the Taylor Lab for generous equipment sharing. Qu lab members are greatly appreciated for insightful discussions and technical support. CPC is supported in part by a Presidential Fellowship of The Ohio State University. This project received partial support from the National Science Foundation (1758912) and the Robert A. Welch Foundation (C-1565 to YJT).

